# Reset Networks: Emergent Topography by Composition of Convolutional Neural Networks

**DOI:** 10.1101/2021.11.19.469308

**Authors:** T. Hannagan

## Abstract

We introduce the Reset model, a composition of neural networks - typically several levels of convolutional neural networks - whose outputs at one level are gathered and reshaped into a spatial input for the next level. We show that units in Reset networks self-organize into clusters when trained on MNIST, Fashion MNIST, CIFAR-10 and CIFAR-100. We then show that a stronger type of self-organization, reminiscent of the topography found for numbers in parietal cortex, arises when number images are mapped onto developmentally realistic number codes. We outline the implications of this model for theories of the cortex and developmental neuroscience.

## Introduction

Convolutional Neural Network (CNN hereafter) classifiers have now been shown beyond any reasonable doubt to predict activity in human visual cortex. However, a very salient aspect of the latter is the clustering, observed in ventral Occipitotemporal cortex (vOTC hereafter), of units selective for faces, houses and other prominent visual categories. These clusters are often called “categorical areas” [1].

Categorical areas cannot be explained within the standard CNN classifier framework, because they respond to high level features in the stimulus, and yet are spatially extended objects in vOTC. On the contrary, CNN classifiers by design trade-off spatial dimensions for feature channels as information is fed-forward. In the deepest layers of a standard CNN classifier, where features are most complex and would have a chance to capture the responses of categorical areas, these features also have little if any spatial arrangement left.

In this article, we show that requiring of CNN outputs to serve as input to other CNNs downstream is sufficient for self-organization to take place. As the input space is in a sense reset, we call these models Reset networks.

## Reset networks

Reset networks are compositions of several neural networks - typically several levels of CNNs - whose outputs at one level are reshaped into a spatial input for the next level. We will show that they implement a sequence of neural spaces where networks performing similar computations end-up being neighbors, as do units that are selective to the same input.

The general form of a Reset network is shown in Figure 1. It has an arbitrary depth of levels, each consisting of several networks operating in parallel on the same input. The next three requirements can be relaxed, but will be followed in the remainder of this article:

**Figure 1.**
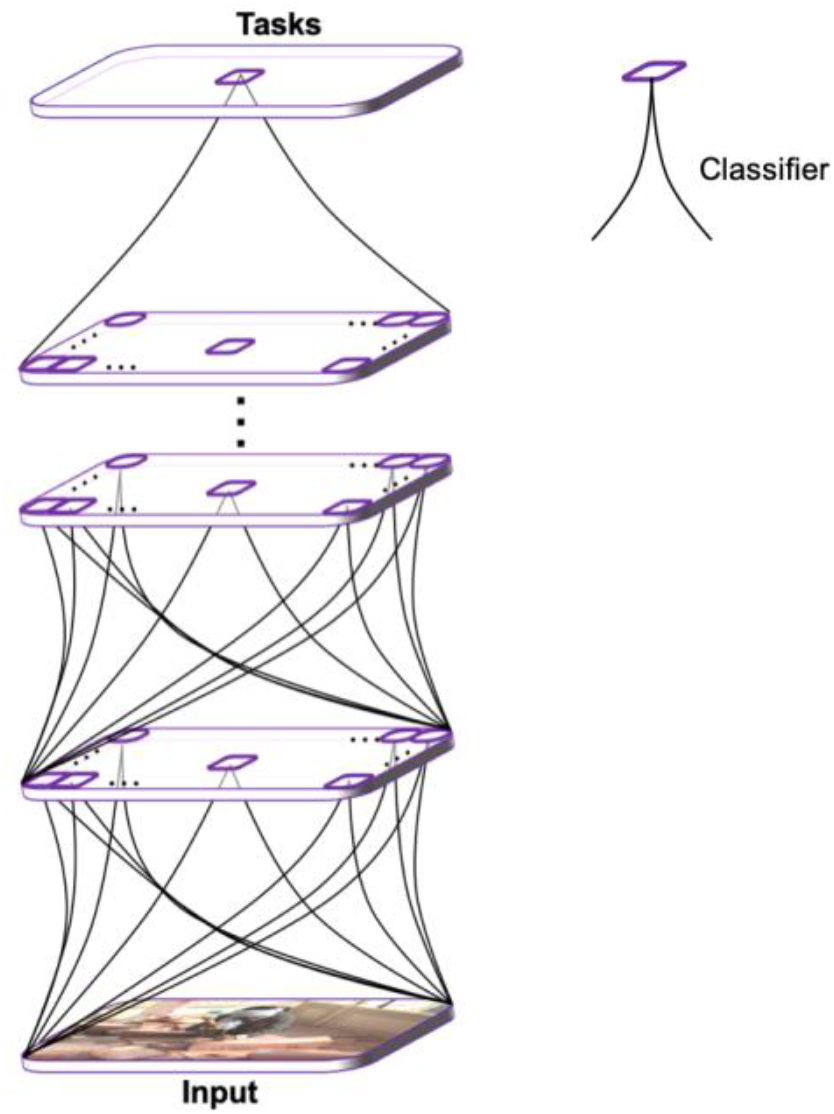
A Reset network is a differentiable neural system with an arbitrary number of levels, where each level itself consists of a spatial arrangement of deep neural networks.

1. At any level, all networks are independent processors: they do not share any weight parameters and do not project to each other laterally.

2. At any level, all networks receive, as a common input, the reshaped and concatenated outputs of all networks from the level below.

3. The last level of the model is the only output level, where error signals for all tasks are received.

Reset networks include in particular the family shown in Figure 2, where level 1 is obtained by reshaping and concatenating the outputs of nxn parallel networks into a single map, called “grid” hereafter, which then serves as input for a final network. We refer to such systems as Reset Networks of depth 2 and width n, or Reset(n), we will sometimes also write Reset(n, m) to further specify the grid’s width m in terms of units.

**Figure 2.**
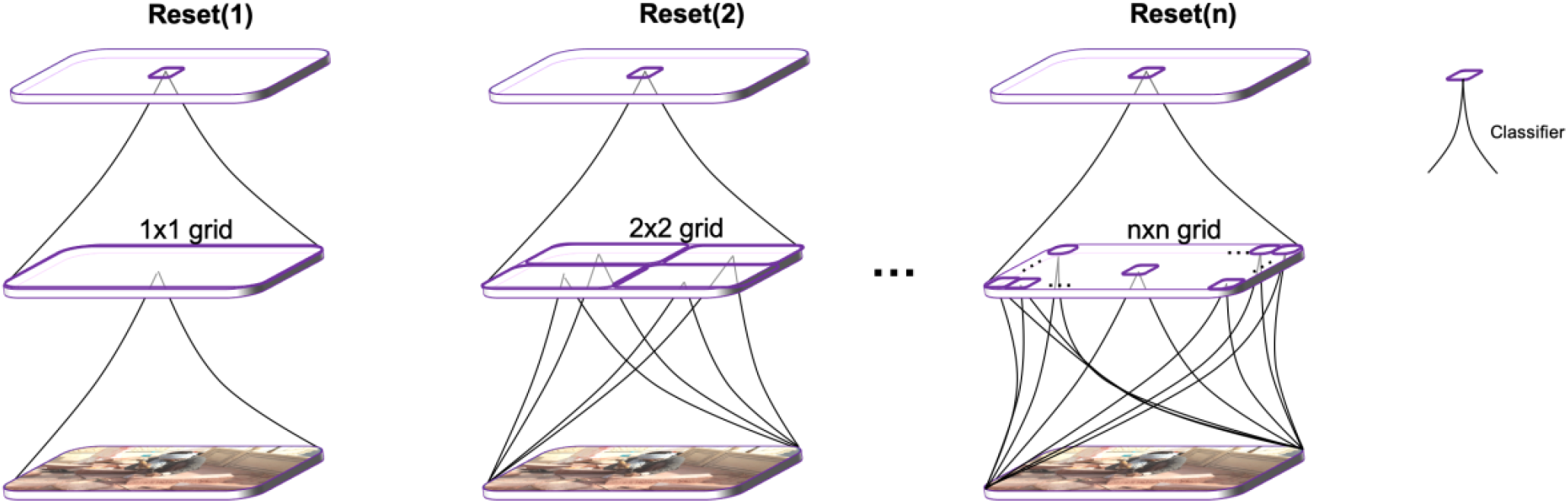
A family of depth 2 Reset networks, with nxn intermediate grids, for increasing n.

We demonstrate that Reset networks can perform classification at scale while also exhibiting emergent topographic organization. Our code is available on GitHub: https://github.com/THANNAGA/Reset-Networks.

## Results

### Reset networks show clustering for MNIST, Fashion MNIST and CIFAR-10

We start by training Reset(8) networks on MNIST, Fashion MNIST and CIFAR-10. In each case, the networks reached standard performance levels on the test sets, but more interestingly, Figure 3 shows the networks’ grids after 20 epochs of training.

**Figure 3.**
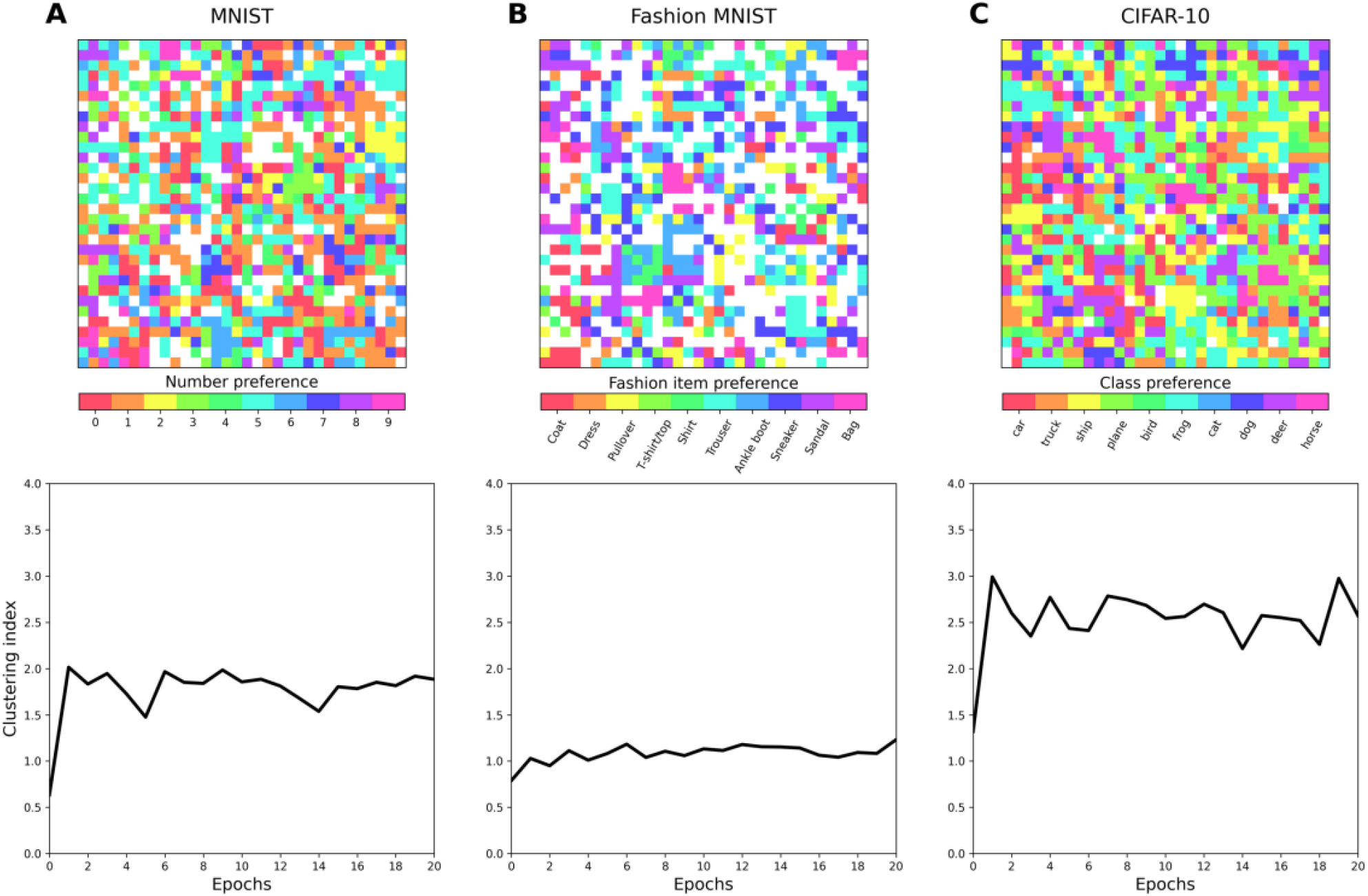
Reset networks with 8×8 grid and depth 2 are trained on MNIST (A), Fashion MNIST (B) and CIFAR-10 (C). (Upper Line) Number preferences on the network grid show topography. (Lower Line) Clustering is defined as the average size, over all target classes, of the connected components on the grid for this class: it is always higher than that obtained for a shuffled map (positive clustering index).

The upper plots in Figure 3 present converged preference maps -the class preference of each unit on the 32×32 grid of the trained model-whereas the lower plots quantify the amount of clustering on each map, at each point during training. A unit’s preference is given by the highest d-prime, over each target class, of the unit’s responses to this target class against all other classes. If this maximum d-prime is below an arbitrary threshold, set to 2 throughout this article, the unit is said to have no preference (white units on the preference map).

We also compute clustering as the average size, over all target classes, of the connected components present on the map for this class. The final clustering index presented Figure 3 is the deviation of clustering from chance: it is obtained by subtracting the clustering measured for shuffled maps to that of the true maps. More details can be found in Supplementary Material (Figure S1).

Figure 3 shows that there is clustering for each of the three domains considered, with some variations across domains: for instance, CIFAR-10 elicits more clustering, while Fashion MNIST produces overall fewer category selective units, and thus less clustering. Figure 3 also shows that clustering happens quite quickly: it is essentially in place after the first training epoch.

One might be forgiven to think that clustering in this model only comes from the concatenation of outputs. To assess whether this is the case, we measure clustering in the family of Reset(n) networks for n = 1, 2, 4 and 8. The grid of Reset(1) has no concatenation, while that of Reset(8) is obtained by concatenating the output units of 8×8=64 subnetworks. We emphasize that although n varies, the size of the grid remains fixed at 32×32 units.

It can be seen on Figure 4 that clustering is always non-zero for all n, and that there is an obvious tendency of clustering to increase with n. Therefore, concatenation accounts for a significant proportion of clustering, but it is not a necessary condition: the Reset(1) curve shows that simply composing two CNN classifiers already will produce clustering.

**Figure 4.**
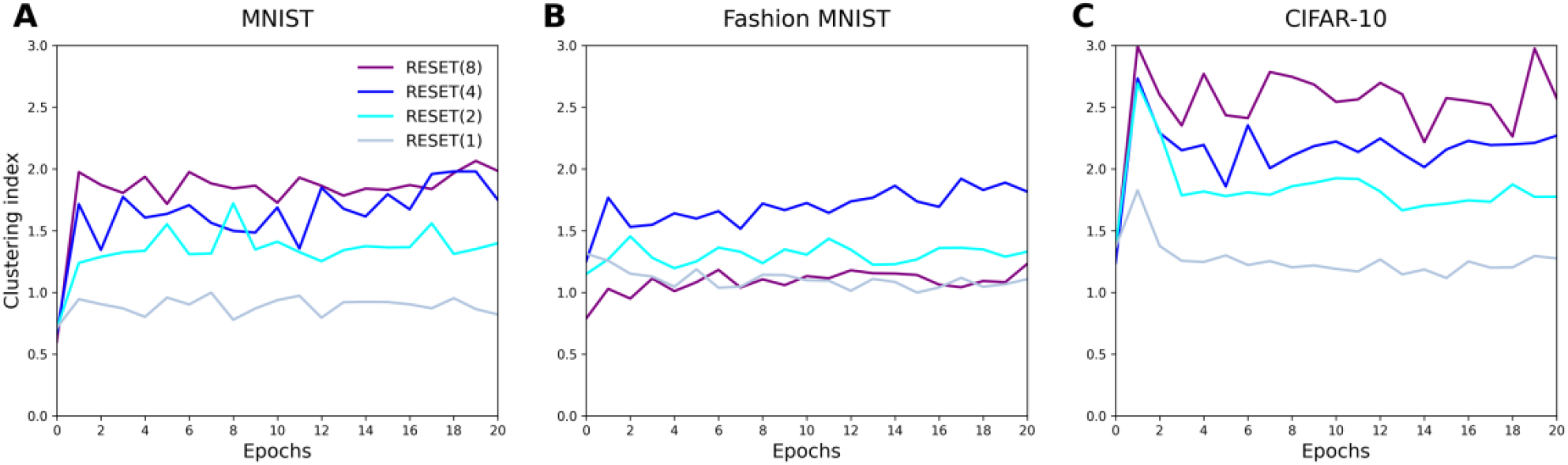
Clustering curves for Reset networks of depth 2 and size 1, 2, 4 and 8 trained on MNIST (A), Fashion MNIST (B) and CIFAR-10 (C). Clustering is non-zero for all networks, and tends to increase with n.

### Reset networks and categorical areas in vOTC

In vOTC, more than two decades of studies have established the presence of areas selective for a few prominent visual categories, in particular faces, body parts, tools, houses, and words. There is no shortage of scalable deep learning models that can reproduce complex responses in the visual system but lack topography, and conversely, topographic models have long been proposed which lack the ability to perform at scale (see for instance [2] for a discussion). To our knowledge, as of 2021 only one model, TDANN [3], can claim both topography and scale at the same time. We return to this model in the discussion, explaining why its treatment of topography is problematic, requiring as it does two disconnected concepts of space to coexist. By contrast, the way Reset networks achieve topography at scale is conceptually straightforward.

The upper line in Figure 5 shows category preferences after training Reset networks of sizes 1, 2, 4 and 8. Only 3 macro-categories are considered – objects, houses and people – which were obtained by aggregating the relevant Cifar-100 classes (see Supplementary Material 1). Though many units have no special preference for these macro-categories, clustering is still obvious on the maps. There also appears to be clustering at the subnetwork level, which often specialize for specific categories. The phenomenon is particularly obvious in the case of Reset(2), as the upper-left subnetwork remains insensitive to any of the 3 categories, the lower right subnetwork has units specializing for all 3 categories, while the lower left and upper right ones specialize for people and objects, respectively. The lower line in Figure 5 also shows that by and large, clustering tends to increase with the number of epochs and with the size of the Reset network.

**Figure 5.**
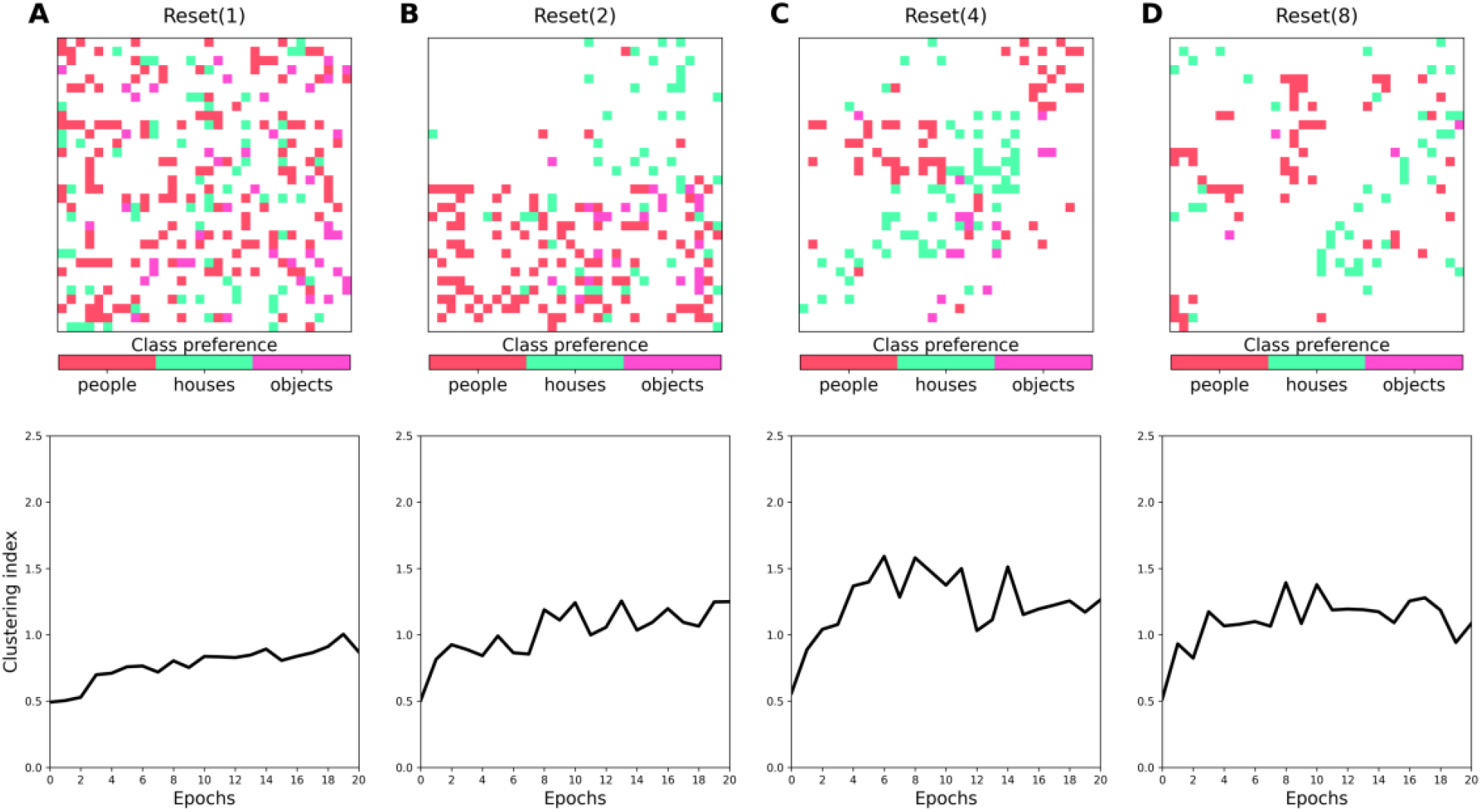
Clustering in Reset networks of depth 2 and sizes 1 (A), 2 (B), 4 (C) and 8 (D) trained on CIFAR-100. (Upper line) Unit preferences on the network grid show clusters for objects, houses and people. (Lower line) As previously observed for other datasets, clustering is non-zero for all networks, and has a tendency to increase with n.

### Reset networks and topography for numbers in parietal cortex

So far we have investigated clustering, which in the context of classification can be defined as the spatial proximity of units that respond to the same class. Though related a notion, clustering is not exactly synonymous with topography.

Cortical topography in the strict sense is the notion that “nearby neurons in the cortex have receptive fields at nearby locations in the world” [4]. However, the term has come to take a wider meaning: it is often understood as applying also to local fields or voxels as well as to neurons, and to refer to any kind of selectivity, not just location selectivity. In this wider sense, topography is a widespread phenomenon in brain imaging, observed throughout the visual cortex as well as in some associative areas.

In parietal cortex, voxels selective for similar numbers are more likely to be contiguous: such a number topography is not yet well understood [5], though some models have provided partial answers [6]. Although we have just seen that Reset networks will self-organize when trained to classify the hand-written digits of MNIST, this task is not entirely satisfying from a developmental and neuroscientific point of view: it is very likely that kids map written digits not onto one-hot labels, but onto pre-existing number representations that have a quite specific format.

The nature of these codes has been studied in [6]: a lot of experimental data could be explained if number codes were sparsely distributed vectors, with a larger overlap between successive number codes as numbers increase. Therefore, it would be more convincing if Reset networks could reproduce number topography by mapping digit images onto these realistic number codes.

As Figure 6 shows, a sequence of Reset(1), (2), (4) and (8) networks -all with the same grid size of 32×32 units-can be trained to map images of digits onto number codes, and succeeds in reproducing topographic organization.

**Figure 6.**
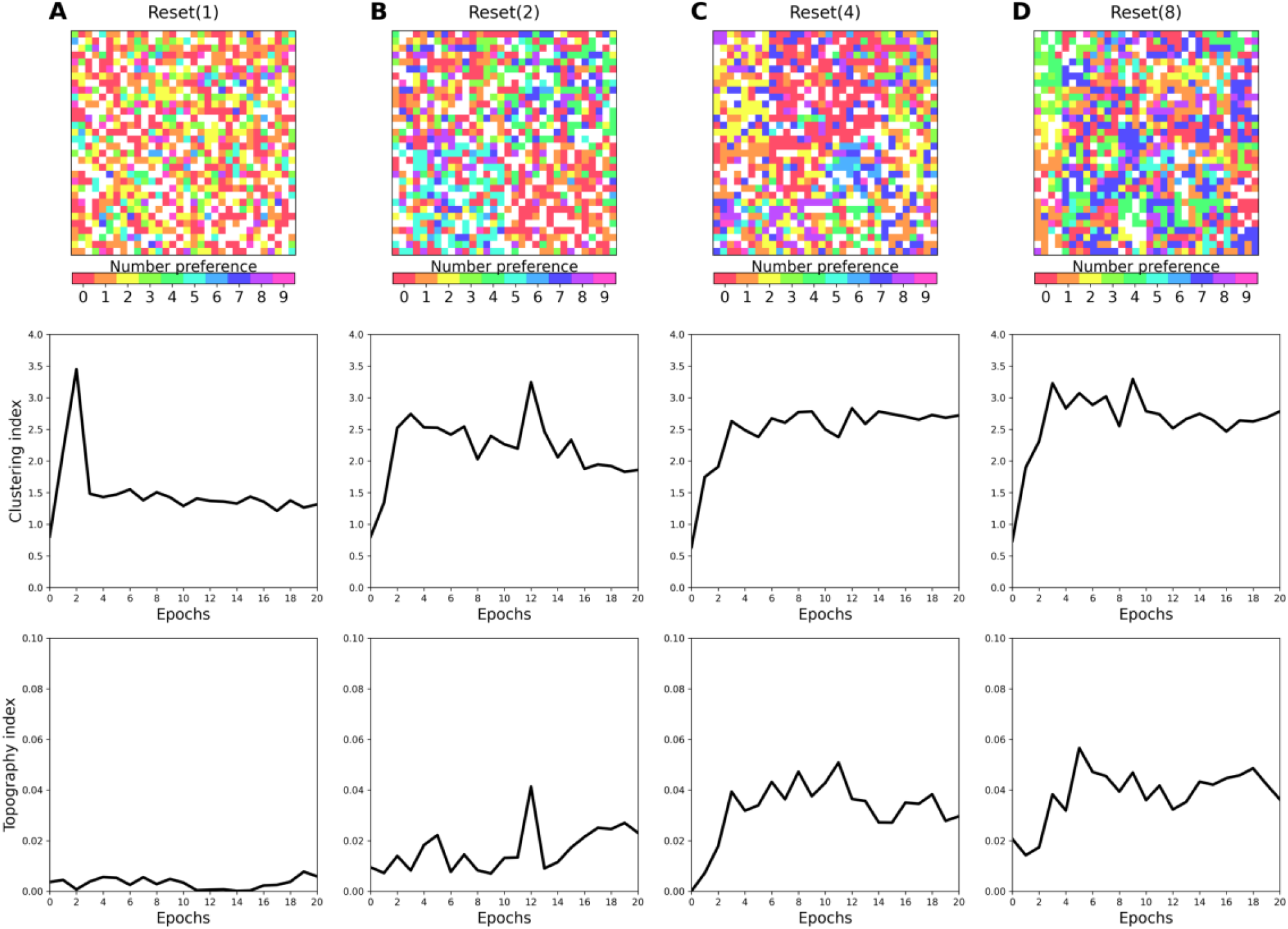
Self-organization for numbers in Reset networks of sizes 1 (A), 2 (B), 4 (C) and 8 (D). (Upper line) Number preferences on the networks’ grids. (Middle line) Clustering takes place in all networks, with a systematic sharp increase early in training, and a tendency to increase with network size. (Bottom line) Topography is close to absent in Reset (1), but otherwise measurable in Reset(2), Reset(4) and Reset(8), increasing with network size and training epochs.

Number topography is visible Figure 6 (upper line) in the maps of number preferences, and is quantified in the clustering curves (middle line), where it can also be seen to emerge quickly during training. Clustering is always quite significantly above what it is for a shuffled selectivity map (positive clustering index). Notably, there is a tendency of subnetworks to specialize for specific numbers, or numbers in the same ballpark.

Consider in this respect the grid of Reset(2), whose lower left quadrant specializes for numbers in the higher range (between 6 and 8), while its lower right quadrant specializes for small numbers like 0 and 1. This type of modularity at the subnetwork level can also be seen in the evolution of reset network preferences during training (see https://github.com/THANNAGA/Reset-Networks/tree/main/Topography%20for%20numbers for time-lapses of grid maturation in Reset(2) and Reset(4)).

Because in this task, the codes onto which number images are mapped have non-degenerate similarities, unlike the binary similarities of one-hot labels used in the previous tasks, one can expect not only clustering, but also topography proper to emerge on the Reset networks’ grids.

We now introduce a topographic index for our preference maps. We define topography as the average, over all units on the grid, of the proximity of a unit’s number preference to those of its 8 immediate neighbors. The topography index presented in bottom line of Figure 6 is then simply the deviation of this topography from chance: it is obtained by subtracting the average topography measured for shuffled maps to that of actual maps (see Supplementary material 1). There is topography as soon as the index is reliably above zero.

The lower plots in Figure 6 show that topography is widespread in Reset networks for this task. With the exception of Reset(1), topography is always much higher than the chance level of 0 (no topography). Two more effects also immediately stand out: topography tends to increase with training, as well as with n -the width of the network’s grid.

Pretrained Reset networks are available at https://github.com/THANNAGA/Reset-Networks/tree/main/Topography%20for%20numbers along with time-lapses showing how number topography evolves during training.

## Discussion

The main insight of Reset networks is that during training, local processing at level L exerts a pressure on the networks’ output from level L-1 to organize in order to solve the task, distributing work in a way that creates clustering or topography. We now discuss some outstanding issues, questions and prospects.

### Classification performance

We have shown that Reset networks can classify standard computer vision datasets such as CIFAR-100. However and as Figure 7 shows, at this stage their performance remains disappointing, only at best matching that of a single Resnet 20, while having many more parameters.

**Figure 7.**
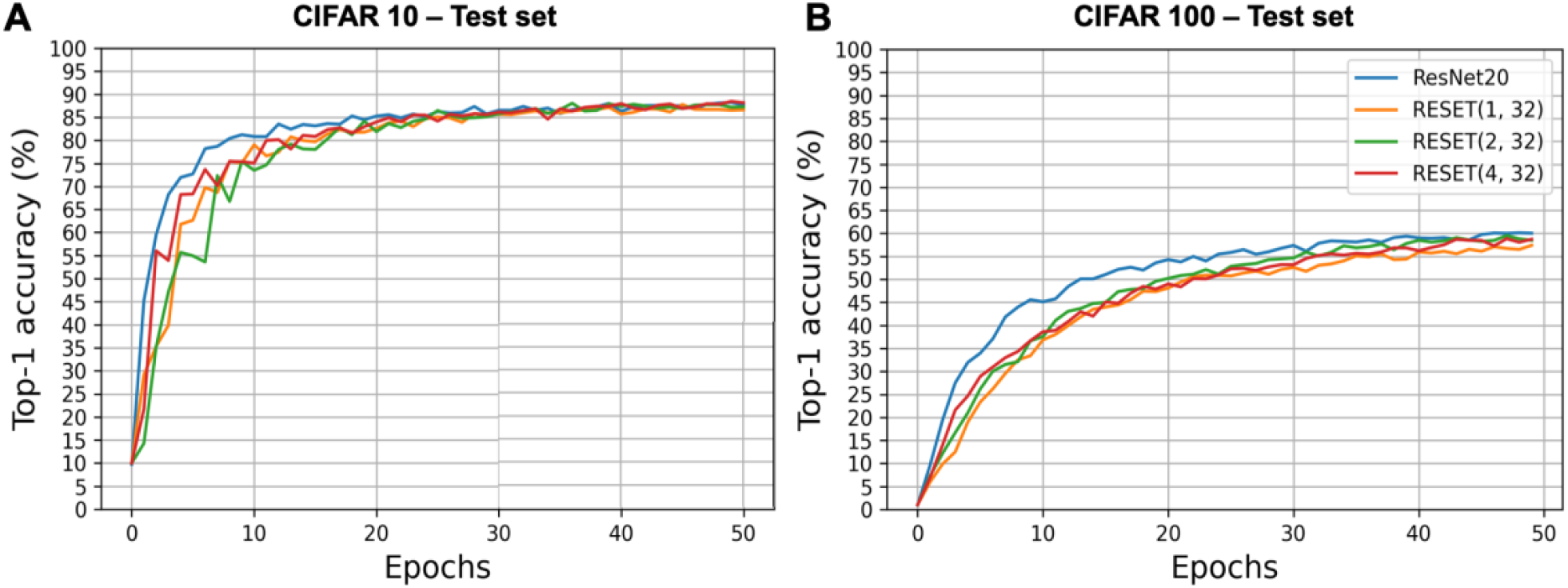
Top-1 accuracy of Reset Networks on CIFAR-10 (A) and CIFAR-100 (B). Composing several CNN classifiers does not destroy classification abilities, though unsatisfyingly, Reset networks can currently at best only converge to the same performance as a single Resnet20. The notation Reset(n, 32) specifies that the outputs of nxn networks are reshaped into a map of 32×32 units.

We also observe that the full resources of the Reset network don’t seem to be used: some subnetwork units are more active than others.

This is not entirely due to the over-parameterization of the networks presented here, as it also happens with smaller subnetworks (unreported simulations), but the issue can be alleviated to some extent by using dropout, or another kind of regularization on the grid.

### Regularization by auto-encoding

In the course of our investigations, we have observed that Reset networks performed much better when the second level had 2 subnetworks: one that classified the input, and another that tried to reconstruct the input from the grid. We trained Reset networks of widths 1, 2 and 4 to classify Cifar-100, with and without adding a subnetwork to reconstruct the input. We then presented 1000 images from the test set to each of the 6 Reset networks, and collected their activations on the grids.

**Figure 7.**
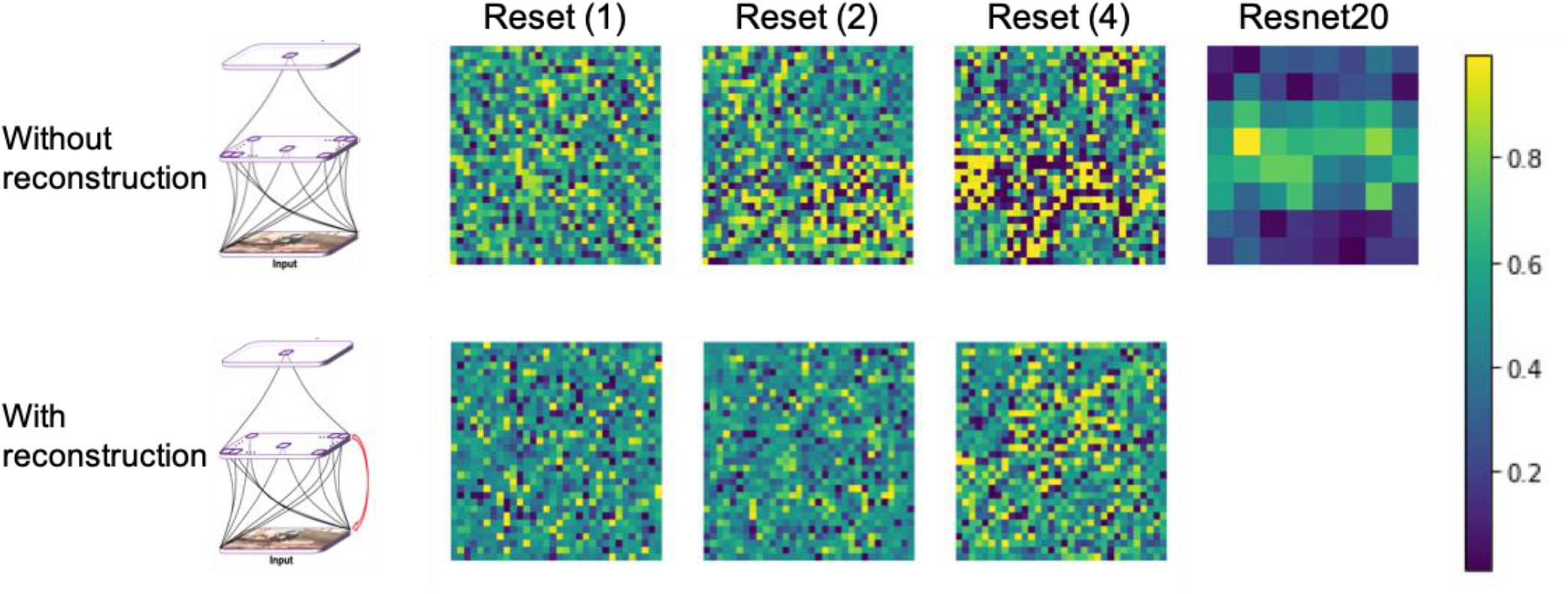
Normalized activation on the grid of Reset networks of width 1, 2 and 4, trained to classify Cifar-100, with and without input reconstruction from the grid. Reconstruction forces the networks to use grid resources more evenly. Activity in the (2d-reshaped) dense layer of Resnet20 is shown for comparison.

Figure 7 (upper line) shows that activation is not evenly distributed on a Reset network’s grid. Units often become polarized, in the sense that many units are either rarely activated (dark purple) or on the contrary very often activated (yellow). This polarization effect becomes stronger within some subnetworks, as network width grows. Figure 7 (lower line) also shows that polarization can be alleviated by introducing a level 2 subnetwork whose task is to reconstruct the input (red arrow on the model’s figure).

Auto-encoding in this situation appears to act as an efficient regularizer for classification, forcing activation to be distributed across the whole grid rather than to be drawn by one, or just a few subnetworks. Such regularization effects of auto-encoding have been reported before for standard classifiers [7]. The novelty in Reset networks is that input reconstruction must be accomplished using the information from the whole grid: this suggests that in visual cortex, some feedback connections between distal cortical areas actually function as regularizers of cortical spaces.

### Topography in Reset networks

Reset networks constitute a novel mechanism for topography to emerge in deep learning. We have presented evidence that Reset networks can reproduce examples of clustering and topographic organization: in parietal cortex, when mapping number images into number codes, and in vOTC, when classifying CIFAR100 images. The fact that self-organization occured for different datasets suggests that it is specific to neither of those, but inherent to the model’s architecture.

Consequently, we would also expect topography, even if a Reset network was trained to classify natural images into number classes. It would also be interesting to find out whether topography for numbers could emerge in an auto-encoding Reset network, in absence of any teaching signal explicitly related to number.

### Alternative to Reset networks

Despite the aforementioned tension between CNNs and vOTC, recent innovative work has shown that categorical areas can indeed be simulated in Topographic Deep Artificial Neural Networks, or TDANNs [3]. In TDANNs, topography is achieved by invoking a separate entity – a “cortical tissue map” – and assigning arbitrary locations on this map to units in the dense layer of the network, before introducing a loss regularizer that penalizes wiring length on the map during training.

Since the mechanism realizing this mapping is unspecified, the ontological status of space in the model is problematic. Two different notions of space exist here that can in principle contradict each other: the spatial coordinates of units in the model’s convolutional feature maps, and the spatial coordinates of the cortical tissue map.

This issue is not brought to the forefront in extant TDANNs, because cortical tissue maps are restricted to the upper dense layers of the model, where locality is lost and units don’t have coordinates. However, there is no reason why cortical tissue maps couldn’t also be invoked for the lower, convolutional levels of the network, with much less interpretability. By contrast, Reset networks achieve topography with a single concept of space.

### Adding networks when necessary: the width and depth of Reset networks

A limit of the current CNN-to-visual-cortex mapping endeavor is that most if not all studies attempt to predict cortical responses from a single deep CNN classifier, trained on a single task. Though understandable, these two simplifications nevertheless make the model qualitatively quite different from the visual system, which is shaped by many different tasks other than classification (e.g. visual tracking, naming), and involves different processing streams.

Reset networks align well with a view of neural development in which, in addition to recycling neural material, new resources can be recruited if needed. Learning a new task could require only to widen the system by adding a network at the current level, with different networks possibly trained on different tasks. If expertise from previously learned tasks is required, the system could be made deeper by reshaping network outputs at the current level and creating a new level. In order to really contribute to continual learning theory, it is now necessary to better specify the mechanisms of network growth within the Reset network approach, that would prevent interference between functions, old and new.

## Conclusion

Reset networks show that deep CNN classifiers can self-organize when they are composed with one another. This finding bears on the twin phenomena of clustering and topography, which pervade the cortex. In this view, the cortex should not be modeled as a single classifier, however deep and richly organized, but as a sequence of levels of neural network classifiers. This in turn rests on the premise that the cortex has the ability to compose networks when necessary, an operation that remains to be observed experimentally.

## Acknowledgments

We thank Thibault Fouqueray for early and stimulating discussions where this idea was seeded, Florence Bouhali and Evelyne Eger for ideas and encouragements on Reset networks. We also thank Hyodong Lee for email exchanges that provided useful information on Topographic Deep Artificial Neural Networks.

## S1. Quantifying clustering and topography

### Clustering index

- Our procedure to quantify clustering is the following:
- Collect average activations for categories A and B on the grid.
- Possibly smooth this activation using a 2d Gaussian kernel (this step was skipped in our analyses).
- Compute the d-prime of A over B for each unit on the grid.
- Threshold the resulting map of d-primes.
- Clustering for a given category is the average number of units in the connected components of this map.

The procedure is illustrated on Figure S1 for the animal vs objects contrast.

**Figure S1.**
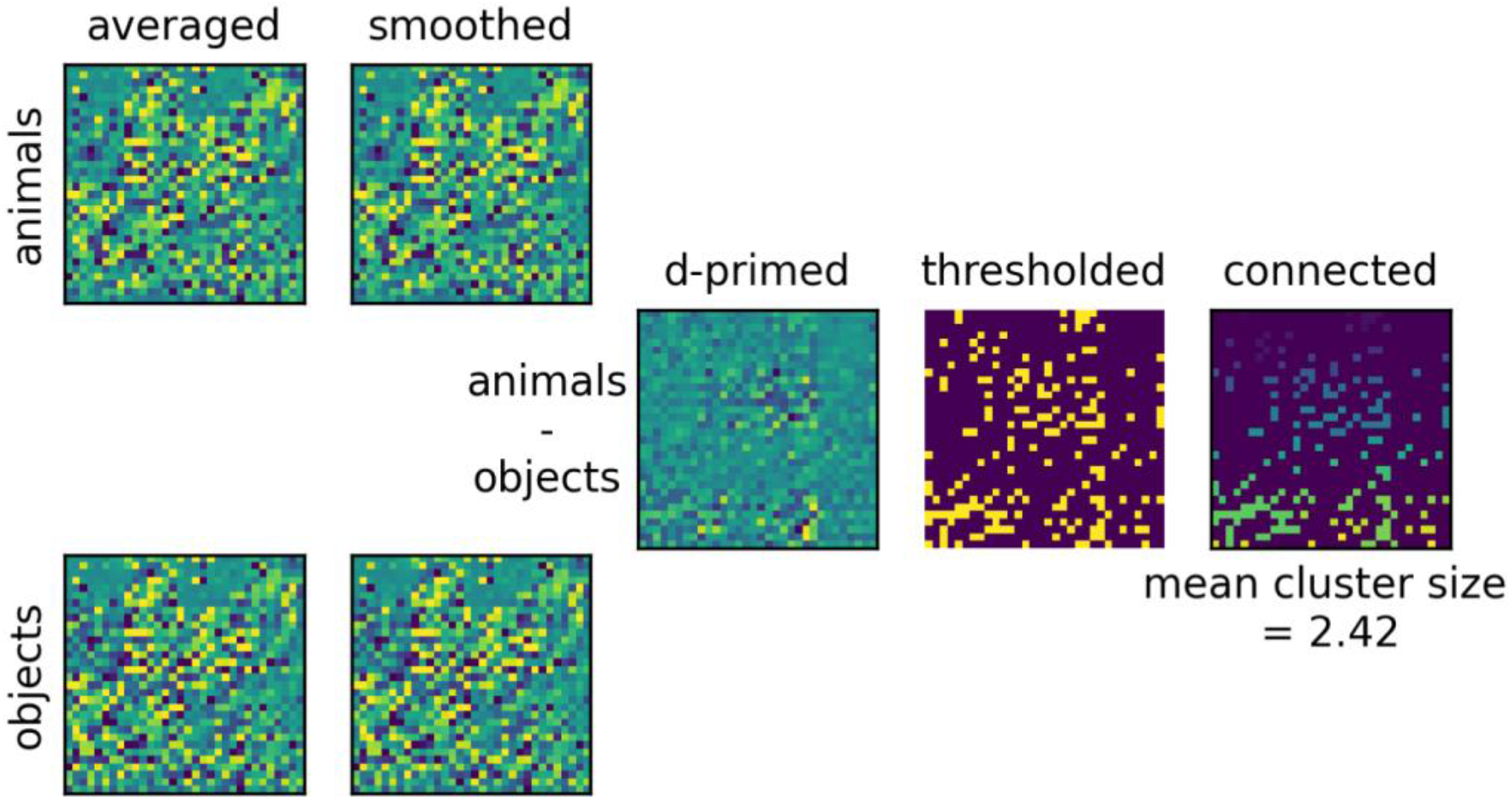
Quantifying clustering using d-primed maps of activations. After thresholding of the map of d-primes, we compute its connected components (color coded in the rightmost plot). Clustering is then defined as the average number of units in the clusters. The divergence of this quantity from that obtained for shuffled preference maps gives our final index.

We call C_M_the mean, over all contrasts of interest, of the clustering measures obtained by the procedure above. We also perform exactly the same computations, averaged over 20 random permutations of M, to obtain a control clustering score for shuffled maps, C_s(M)_. Our final topographic index is then given by |C_M_−C_s(M)_|.

### Topography index

Consider a unit x and its neighborhood v(x) on a preference map M over n classes. Define *p*(*x*) as the class preference of unit x, and <.> as the average operator. The topography of map M, noted *T*_*M*_, is given by the average over x of the similarity in preferences between x and all units in v(x):

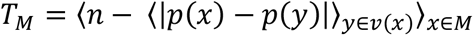

The control quantity *T*_*s*(*M*)_ is obtained in the same way, but further averaged over 20 random permutations of M. Our final topographic index is then given by |*T*_*M*_ −*T*_*s*(*M*)_|.

*Definition of test classes for the clustering simulations on CIFAR-100*

**Table S1.**
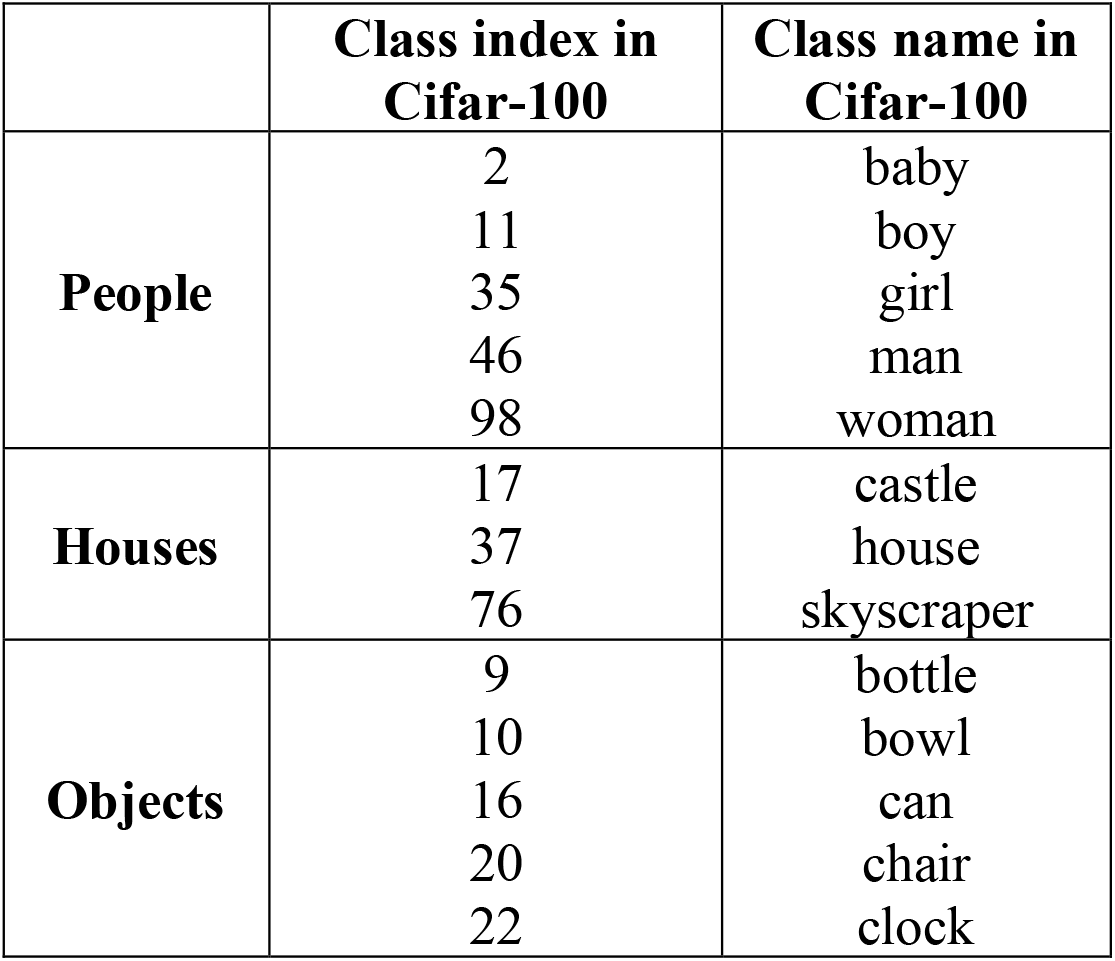
Macro-classes defined from CIFAR100 and used during test.

|  | **Class index in**<br>**Cifar-100** | **Class name in**<br>**Cifar-100** |
| --- | --- | --- |
| **People** | 2 | baby |
|  | 11 | boy |
|  | 35 | girl |
|  | 46 | man |
|  | 98 | woman |
| **Houses** | 17 | castle |
|  | 37 | house |
|  | 76 | skyscraper |
| **Objects** | 9 | bottle |
|  | 10 | bowl |
|  | 16 | can |
|  | 20 | chair |
|  | 22 | clock |

## S2. Modeling Number topography

### Dataset

Our custom-made dataset comprised 6000 exemplars of number images between 0 and 9, paired with as many number codes from [4]. Number images were black on white 32×32 pixel images, in 9 possible fonts (arial, lato, openSans, ostrich, oswald, PTN57F, raleway, roboto and tahoma), 6 x-locations and 24 y-locations. The dataset is available as numpy arrays at https://github.com/THANNAGA/Reset-Networks/blob/main/Topography%20for%20numbers/dataset_number_topography.zip

The 10 number codes onto which those images were classified were 100 dimensional vectors taken from [3], obtained by power iteration of a randomly and locally connected matrix.

**Figure S2.**
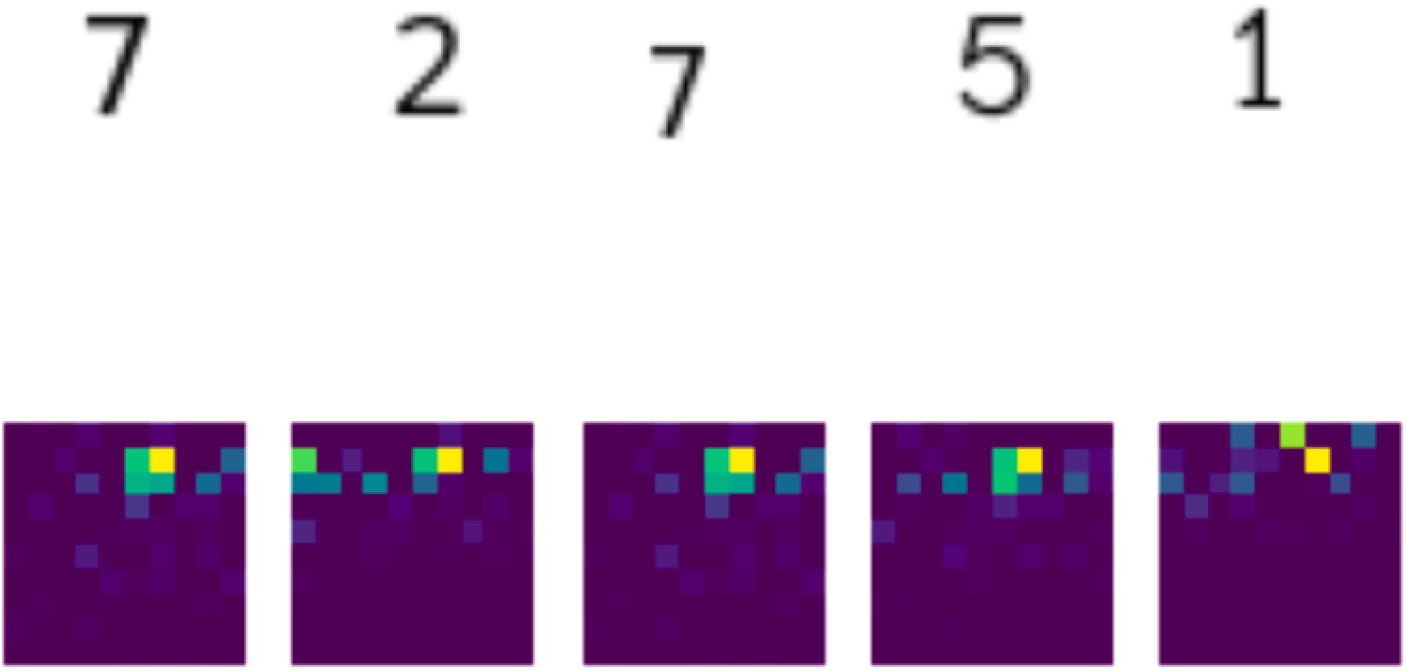
Input and labels of the number dataset used for our number topography

These number codes are sparse, overlapping and real-valued: this implies that the networks trained on this dataset realize a multi-label classification with soft labels.

### Models and training

The networks had the following number of parameters:

**Table S3.** Number of parameters in the networks trained on the number task.

|  | # Parameters |
| --- | --- |
| **Reset(1)** | 610918 |
| **Reset(2)** | 1418134 |
| **Reset(4)** | 4646998 |
| **Reset(8)** | 17562454 |
| **Resnet20** | 275572 |

All models were trained for 20 epochs using a Binary Cross-Entropy loss, the Adam optimizer with a learning rate of 0.001, and a dropout of 0.5 applied to the grid.

